# A Chlorophyte alga utilizes alternative electron transport for primary photoprotection

**DOI:** 10.1101/2020.03.26.010397

**Authors:** Maxwell A. Ware, Darcy Hunstiger, Michael Cantrell, Graham Peers

**Affiliations:** Department of Biology, Colorado State University, Fort Collins, Colorado 80523

## Abstract

*Desmodesmus armatus* is an emerging biofuel platform producing high amount of lipids and biomass in mass culture. We observed *D. armatus* in light-limiting, excess light and sinusoidal light environments to investigate its photoacclimation behaviors and the mechanisms by which it dissipates excess energy. Chlorophyll *a:b* ratios and the functional absorption cross section of photosystem II (PSII) suggested a constitutively small light harvesting antenna size relative to other green algae. *In situ* and *ex situ* measurements of photo-physiology revealed that nonphotochemical quenching (NPQ) is not a significant contributor to photoprotection, but cells do not suffer substantial photoinhibition despite its near absence. We performed membrane inlet mass spectrometry analysis to show that *D. armatus* has a very high capacity for alternative electron transport (AET) measured as light dependent oxygen consumption. Up to 90% of electrons generated at PSII can be dissipated by AET in a water-water cycle during growth in rapidly fluctuating light environments like those found in industrial scale photobioreactors. This work highlights the diversity of photoprotective mechanisms shown in algal systems, that NPQ is not necessarily required for effective photoprotection in some algae and suggests that engineering AET may be an attractive target for increasing biomass productivity of some strains.

**One-sentence summary:** Constitutive small antennae, alternative electron transport and an efficient photosystem II turnover capacity enable *D. armatus* to photosynthesize efficiently.

## Introduction

Photoprotective processes allow the reaction oxygenic photosynthesis to maintain relatively high efficiencies in fluctuating light. The mechanistic understanding of photoprotective processes in algae are mostly based on a limited number of model organisms, such as *Chlamydomonas reinhardtii, Phaeodactylum tricornutum*, as well as inference from observations in plants (e.g. *Arabidopsis, Nicotiana, Triticum*). These species predominantly rely on non-photochemical quenching (NPQ) to dissipate excess energy (Ruban, 2012). This process becomes important for avoiding damage to the photosynthetic system when absorbed light energy cannot be fully utilized by the light harvesting reactions. NPQ is triggered by a ΔpH gradient across the thylakoid membrane that leads to conformational changes in the light harvesting antennae associated with PSII (Horton and Ruban, 1992; Goral *et al*., 2012; Nicol *et al*., 2019).

Algae are a polyphyletic grouping of organisms and this is reflected in the varied light harvesting antennae pigment-protein complexes utilized by algae, despite their similarities in the structure of PSII core proteins (Tokutsu and Minigawa, 2013; Pi *et al*., 2019). Additionally, different processes may fulfil the requirements for photoprotection in different taxa. For example, Streptophyte plants and the model Chlorophyte alga *C. reinhardtii* use the violaxanthin/antheraxanthin/zeaxanthin (VAZ) cycle to assist in the formation of NPQ. However, Stramenopile algae utilize a novel set of related carotenoids for the diadinoxanthin/diatoxanthin cycle for the same purpose (Demers *et al*., 1991). It is also possible that other photoprotective processes significantly contribute to photoprotection in algae such as alternative electron transport (AET; Houille-Vernes *et al*., 2011), photosystem II repair (Key *et al*., 2000) and photorespiration (Niyogi, 2000).

*Desmodesmus armatus* (strain SE 00107; previously referred to as *Scenedesmus armatus)* is an emerging alga of biotechnological interest. It has been demonstrated to detoxify wastewater and has resistance to high metal toxicity (Rugnini *et al*., 2016; Rugnini *et al*., 2017). *Desmodesmus* species have been shown to survive temperature fluctuations of 45°C for 24 h, with less than 13% heat related mortality, and to flocculate readily inside ~2.5 h of settling (Pan *et al*., 2011; Chen *et al*., 2020). *D. armatus* naturally produces commercially valuable nutraceuticals such as lutein and lycopene and can accumulate lipids up to 50% of their dry weight biomass (Pan *et al*., 2011). All these factors contribute to its potential for deployment at a commercial scale.

Although *D. armatus* and other *Desmodesmus* species have been explored as a potential source for biomass, bioproducts and biofuels (Sijil *et al*., 2019; Zhang *et al*., 2016), detailed photosynthetic characterization of this organism has not been performed. *D. armatus* possesses chlorophylls *a* and *b*, the VAZ cycle carotenoids (Tukaj *et al*., 2003). In the closely related species *Scenedesmus quadricauda* NPQ is inhibited by the addition of dithiothreitol, suggesting this group likely utilizes a violaxanthin de-epoxidase mediated NPQ (Masojìdek *et al*., 1999).

There are other notable aspects of photophysiology that can minimize the over-excitation of PSII and maintain a maximum yield of PSII (ΦPSII) in excess light. Changes in the ratios of antennae proteins to reaction center core proteins, and thus pigment ratios, have been demonstrated in plants, *Chlamydomonas*, diatoms and cyanobacteria in response to growth phase, culture/canopy densities and light acclimation history (Cantrell and Peers, 2017; Murchie and Horton 1997; Schagerl and Müller, 2006). Lower absorption cross sections simply capture less light per reaction center and reduce the probability of photodamage in excess light.

When algal blooms or mass cultures reach high cell densities, light limitation can occur. This leads to an increase in pigment per cell concentrations and in antennae proteins compared to core proteins (Neale and Melis, 1986). This ensures a higher probability of light capture. However, for productivity of mass culture, wasteful processes constitute huge losses to productivity (Andersson *et al*. 2019; Melis, 2009). Pigment reduction or truncated light harvesting antennae proteins have been demonstrated to improve solar conversion efficiencies of mass cultures, whilst simultaneously reducing NPQ capacity and rates of photoinhibition in individual cells (Nakajima and Ueda, 1997; Melis, 2009). This was demonstrated in the truncated light-harvesting chlorophyll antenna size *(tla1)* and in LHC translator repressor (*T7*) *Chlamydomonas reinhardtii* mutants (Polle *et al*., 2003, Beckmann J *et al*., 2009).

AET is a collection of photoprotective mechanisms that are gaining greater prominence in algal photobiology research. AET is involved in light-independent and light-dependent balancing of reductant. These processes utilize different forms of reductant to reduce oxygen and produce water. Loss of AET proteins have been demonstrated to reduce photochemical quantum yields and growth rates in *Chlamydomonas* under intermittent light conditions (Chaux et al., 2017; Nawrocki *et al*., 2019). Allahverdiyeva et al. (2013) as well as Andersson (*et al*. 2019) have also shown AET to be an important photoprotective mechanism in cyanobacteria. Importantly, modifying the accumulation of AET-related proteins can improve photosynthetic efficiency and decrease photoinhibition in plants and cyanobacteria (Hasunuma *et al*., 2014; Yamamoto *et al*., 2016; Gómez *et al*., 2018). Together this work suggests a prominent role for AET in photosynthesis, and its application in bioengineering.

The efficient repair of excess-light induced damages to the PSII reaction center D1 protein is another mechanism to mitigate losses associated with photoinhibition (Key *et al*., 2010). PSII turnover occurs at all light intensities, and efficient repair rates are required to maintain a functional PSII reaction center when photosynthesis and photoprotective mechanisms are unable to utilize all energy absorbed by photosynthetic pigments (Tyystjärvi and Aro, 1996). Fast D1 protein repair rates can help maintain a high ΦPSII but cannot operate on the 10^−15^-10^−10^ s timescales of photon capture and transfer that is required to minimize reactive oxygen species (ROS) formation (Ruban, 2012). There are also energetic costs required with the removal of damaged D1 protein and new protein synthesis (Silva *et al*., 2013). Thus, PSII repair is not viewed as a classical photoprotective mechanism, but it is an essential process for maintaining maximum yields of photosynthesis and avoiding sustained photoinhibition.

Bioengineering of light harvesting and photoprotective mechanisms have been demonstrated to yield 15% increased biomass in plants (Kromdijk *et al*., 2016) and 28% in cyanobacteria (Peers, 2015). Engineering targets are however likely to be species and cultivation scenario specific. The focus of this research is to better understand how *D. armatus* adapts to changing light. We sought to investigate how this organism balances light harvesting and energy dissipation between two major dissipative pathways (AET and NPQ), and what effect this has on ΦPSII and growth rates under light limiting, excess light and simulated natural light conditions. Here we show that *D. armatus* has a relatively small functional PSII antennae, high photoprotection capacities and maintains high photochemical quantum yields throughout exponential growth phase. We elucidated the main photoprotective mechanism to be AET, despite the capacity to form limited NPQ. Furthermore, *D. armatus* has a very efficient D1 repair rate which contributes to a high ΦPSII in excess light conditions.

## Results

### Growth in square-wave and sinusoidal light conditions

A preliminary set of experiments set out to characterize the growth stages of *Desmodesmus armatus* (SE 00107) in several light conditions. Cultures were grown under 12:12 h D:N high (HL, 600 μmol photons m^−2^ s^−1^), low (LL, 60 μmol photons m^−2^ s^−1^) and sinusoidal light conditions (0-2,000 μmol photons m^−2^ s^−1^; Fig S1). In all conditions, maximum growth rates were observed during the first four days, coinciding with the highest maximum PSII photochemical yield (Fv/Fm) values and highest average chl *a/b* ratios (ranging from 6-7; Supplemental Table S1). This informed us that the exponential growth phase was within 1×10^5^-1.5×10^6^ cells/ml, and all subsequent experiments were performed during this time.

Fv/Fm measurements show that LL-acclimated cultures maintain a high PSII quantum yield throughout the day without experiencing photoinhibition (Fig 1A). HL-acclimated cultures were photoinhibited during the first 30 min of illumination. This difference in maximum photochemical quantum yield was significantly lower than all other time points (Zeitgeber time, ZT; RM-ANOVA, p<0.001; Tukey’s test, p≤0.013). The only other significant difference in Fv/Fm was at ZT 9 h to 3 and 12 h (Tukey’s test, p≤0.019). Fv/Fm values of HL adapted cultures are not otherwise significantly different between 3 and 24 ZT h.

SL-acclimated cultures had a higher maximum photochemical quantum yields than HL acclimated cultures at every time point during the day except ZT 9 h (Fig 1a; Tukey’s test, p≤0.014). SL conditioned cultures only had significantly higher maximum quantum yields than LL acclimated cultures at ZT 0 and 24 h (Tukey’s test, p≤0.043).

**Figure 1.**
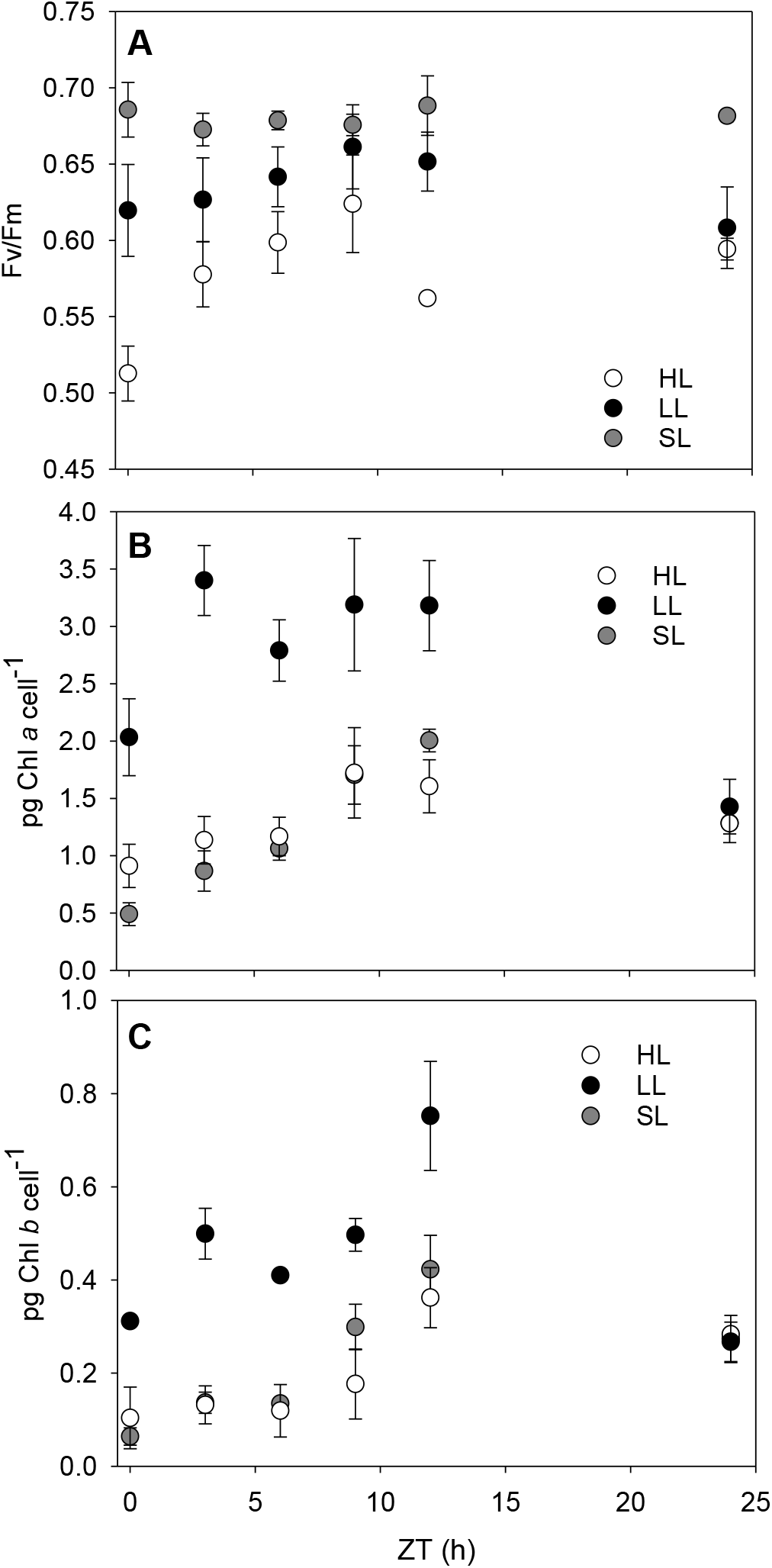
Estimations of PSII maximum photochemical quantum yields (Fv/Fm) and chlorophyll pigment accumulation during a diurnal light cycle. HL and LL correspond to growth with incident light fluxes of 600 μmol photons m^−2^ s^−1^ and 60 μmol photons m^−2^ s^−1^, respectively. SL indicates cultures were grown with sinusoidal daily irradiance in an ePBR photobioreactor. A) *In situ* average maximum photochemical quantum yield of PSII. B) pg chl *a* and C) chl *b* per cell extracted in methanol and calculated spectrophotometrically. Error bars represent standard deviation (n=3).

### Pigment Content

The amount of chl *a* and *b* per cell increased between dawn and dusk in both diurnal square-wave light acclimation histories before declining during darkness (Figs 1B and C). LL-acclimated cells also had significantly higher chl *a* (RM-ANOVA, p≤0.041; Tukey’s test p≤0.08) and chl *b* (RM-ANOVA, p≤0.022; Tukey’s test, p≤0.049) quantities per cell at every timepoint during the photoperiod. Additionally, the chl *a/b* ratio was dynamic during the day in all conditions. Illustrating this, SL acclimated cells had a chl *a/b* ratio of 7.92±1.02 at solar noon before decreasing to 4.83±0.74 (Fig S2).

SL-adapted cells had a biphasic increase in chl *a* and *b* concentrations per cell throughout the daylight period (Figs 1B and C). Gradual increases in pigments were measured from ZT 0-6 h, with a large increase after solar noon. This strategy most closely mimicked HL-acclimated cells.

### Estimations of PSII supercomplex functional size in Fm’ and Fm states

Measurements of functional antenna size in the dark-adapted state vs. in the light adapted state (*in situ)* suggested there was very little physiological adjustment of PSII size throughout the different growth conditions (Fig 2A). LL acclimated cells had a slightly smaller average σPSII than HL-acclimated cultures (Tukey’s test, p<0.05). We tested to see whether this was possibly due to a state transition-like response where light harvesting complexes migrate from one photosystem to the other. However, were unable to observe state-transitions in the species (Fig S3). A slight peak at ~695 nm in the 77K fluorescence emission spectra did suggest that there are some antennae proteins functionally uncoupled from PSII (Fig S3B). Comparing *in situ* ΦPSII to dark-adapted Fv/Fm illustrated minimal downregulation of PSII efficiency due to photoinhibition or NPQ induction in both static-light acclimated conditions (Fig 2B). Despite little observed NPQ, the parameter Fo/σPSII shows no significant changes in the relative abundance of open PSII reaction centers throughout the day (Fig 2C, RM-ANOVA p>0.1). Sinusoidal light cultures did not display a notable deviation from these patterns.

**Figure 2.**
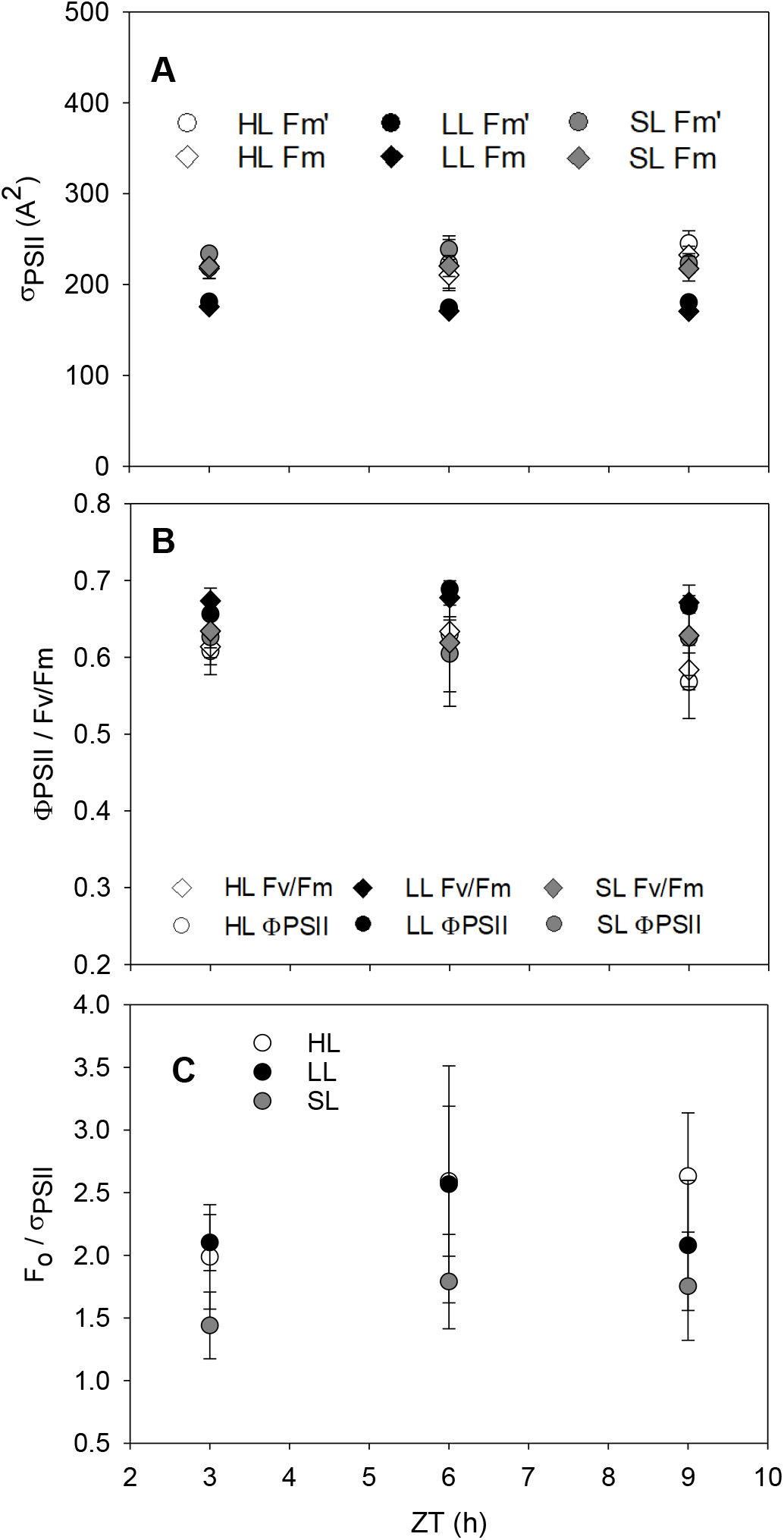
Estimations of PSII functional absorption cross section (σPSII), *in situ* PSII downregulation and concentration of open PSII reaction centers. HL and LL correspond to growth with incident light fluxes of 600 μmol photons m^−2^ s^−1^ and 60 μmol photons m^−2^ s^−1^, respectively. SL indicates cultures were grown with sinusoidal daily irradiance in an ePBR photobioreactor. The *in situ* and dark adapted functional PSII absorption cross section (σPSII) (A) was simultaneously measured with the *in situ* ΦPSII and dark-adapted Fv/Fm (B). Data denoted as Fm’ correspond to ΦPSII, and Fm to Fv/Fm in (A). C) Estimation of functional photosystem II reaction centres (Fo/σPSII) in the Fm state. Error bars represent standard deviation (n=3).

### Chlorophyll fluorescence dynamics

We tested if NPQ is present but not activated *in situ*, or if other photoprotective mechanisms are employed by utilizing photosynthetic irradiance (P-I) protocols. These protocols allow us to simultaneously observe chlorophyll fluorescence kinetics along with photosynthetic gas exchange using membrane inlet mass spectrometry (MIMS). Chlorophyll fluorescence data showed that *D. armatus* does not exhibit a high NPQ capacity, no matter the acclimation history (Fig 3A). LL acclimated cells did have the greatest NPQ capacity, with significantly higher NPQ than HL-acclimated cultures at 0-470 μmol photons m^−2^ s^−1^ (RM-ANOVA, p≤0.019; Tukey’s test, p≤0.022) and at light fluxes greater than 2709 μmol photons m^−2^ s^−1^ (RM-ANOVA, p≤0.038 and Tukey’s test, p<0.031). Estimations of excitation pressure (1-qL) show that despite higher average total NPQ capacities, LL cells were not as effective at stopping the overreduction of Q_A_ (Fig 3B). Indeed, differences in the 1-qL parameter show significant differences between LL and HL conditioned cells at all light intensities (RM-ANOVA, p≤0.007; Tukey’s test, p≤0.008). At close to maximum incident sunlight intensities (~2,000 μmol photons m^−2^ s^−1^), less than 60% of primary electron acceptors are in the reduced state in HL cultures (HL 3B).

**Figure 3.**
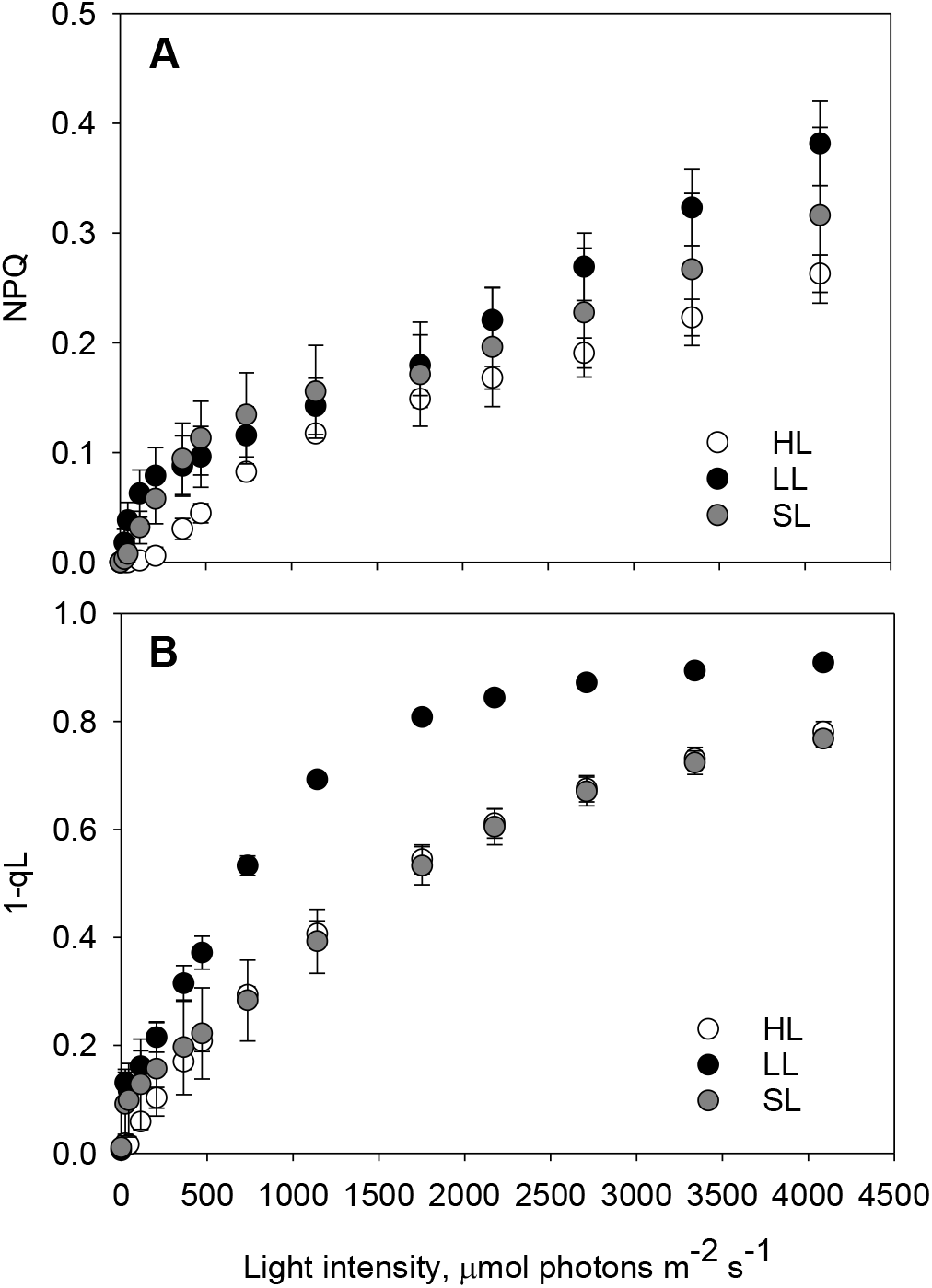
Redox state of the plastoquinone pool (1-qL) and Non-Photochemical Quenching (NPQ) capacities of *D. armatus* cultures measured during an *ex situ* P-I curve. HL and LL correspond to growth with incident light fluxes of 600 μmol photons m^−2^ s^−1^ and 60 μmol photons m^−2^ s^−1^, respectively. SL indicates cultures were grown with sinusoidal daily irradiance in an ePBR photobioreactor. NPQ (A) and 1-qL (B) were measured with a DUAL-PAM fluorometer. 2 μg chl *a* was loaded in 1.4 ml total cuvette volume. Error bars represent standard deviation (n=4).

SL-acclimated cells had significantly lower NPQ capacity than LL-adapted cultures 208 μmol photons m^−2^ s^−1^ (Fig 3A; Tukey’s test, p≤0.043), and higher NPQ capacity than HL acclimated cells at 44-472 μmol photons m^−2^ s^−1^ (Fig 3A; Tukey’s test, p≤0.047). Similarly, to HL-grown cells, SL-grown cells required light intensities of >2,000 μmol photons m^−2^ s^−1^ to reduce most electron carriers (Fig 3B).

In a separate set of experiments, we also exposed cultures to conditions known to induce a large NPQ capacity in other green algae. SL-acclimated cells were grown into late stationary phase (chl *a/b* was <3.0). These were then assayed for NPQ capacity following 5 min of 750 μmol photons m^−2^ s^−1^ of actinic light. These cells were then also treated with UV-B light as an additional method to induce NPQ capacity (Allorent *et al*., 2016; Fig S4). However, neither the stationary cultures nor stationary + UV treated samples displayed significant NPQ *ex situ*, but UV treatment did cause a decline in Fv/Fm.

### Membrane inlet mass spectrometry (MIMS)

We measured gas exchange using MIMS with exogenously added ^18^O_2_, which allows for the observation of both oxygen evolution and consumption reactions in the light. Measurements of gross oxygen evolution show that the photosynthetic capacity of cells depended on acclimation history, with significant differences between conditions at every light intensity 208< μmol photons m^−2^ s^−1^ (Fig 4A, RM-ANOVA, p≤0.027). HL-acclimated cells had significantly higher gross oxygen evolution rates normalized to chl *a*, than LL (208< μmol photons m^−2^ s^−1^; Tukey’s test, p≤0.025) or SL-acclimated cultures (363< μmol photons m^−2^ s^−1^; Tukey’s test, p≤0.045). Oxygen consumption rates did not change significantly during the lowest light intervals (≤363 μmol photons m^−2^ s^−1^), but once the light intensity reached ~470 μmol photons m^−2^ s^−1^, there was an increase in average oxygen consumption at each actinic light interval (Fig 4B). HL-adapted cells have, on average, 3.4x higher net photosynthesis than LL cells, which illustrates the huge adaptation response in this organism. Significant differences between LL and HL cultures in the minimum saturating light intensity (Pmax; Tukey’s test, p<0.001), maximum rates of oxygen evolution (I_K_; Tukey’s test, p<0.001) were also observed (Fig. 5).

**Fig 4.**
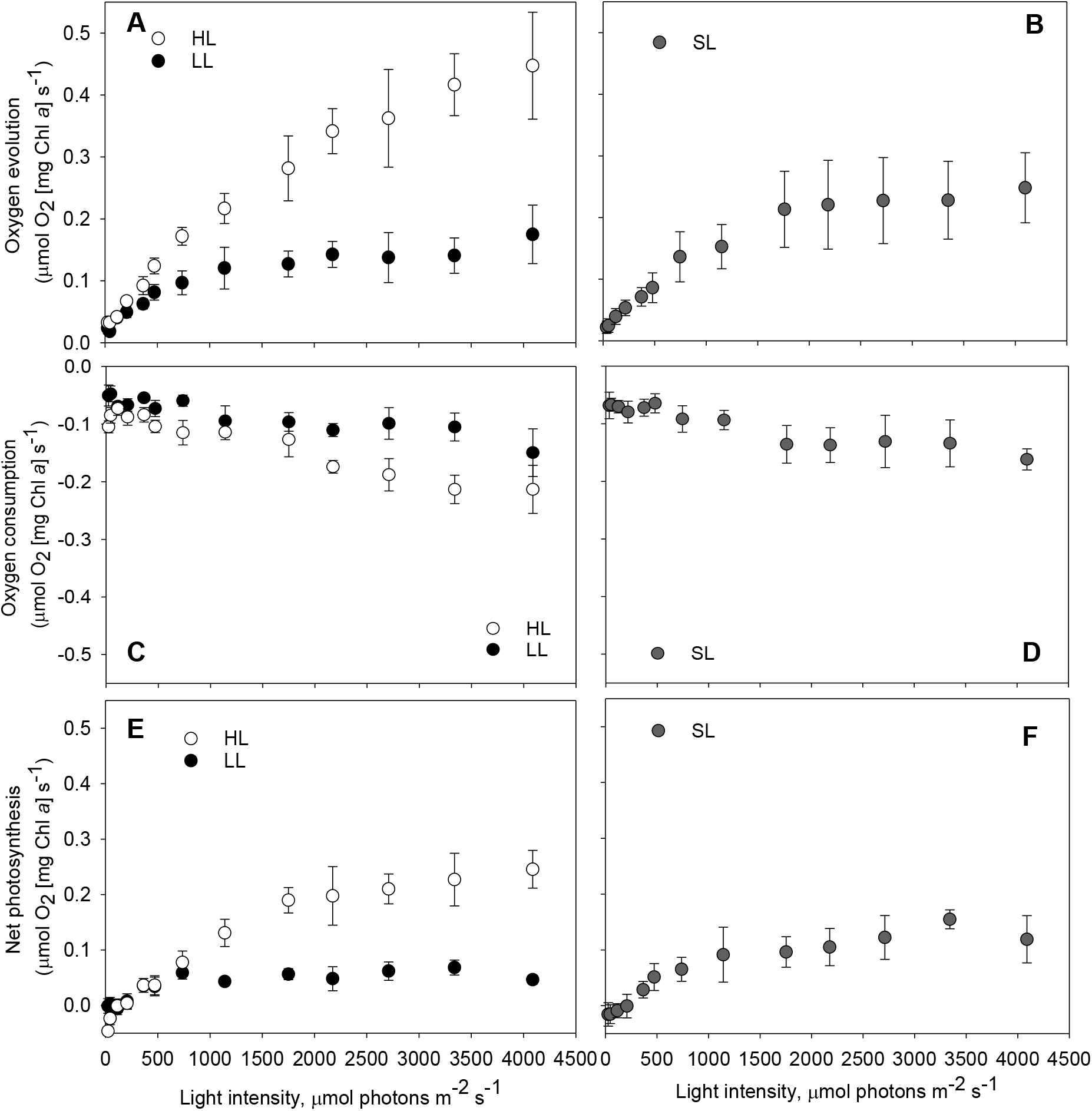
*Ex situ* estimations of oxygen evolution, consumption and net photosynthesis capacity. HL and LL correspond to growth with incident light fluxes of 600 μmol photons m^2^ s^−1^ and 60 μmol photons m^−2^ s^−1^, respectively. SL indicates cultures were grown with sinusoidal daily irradiance in an ePBR photobioreactor. Data were gathered simultaneously with that presented in Figure 3. (A, B) Gross oxygen evolution, (C, D) gross oxygen consumption and (E, F) net photosynthesis was measured using MIMS. 2 μg chl a was loaded in 1.4 ml total cuvette volume. Error bars represent standard deviation (n=4).

**Figure 5.**
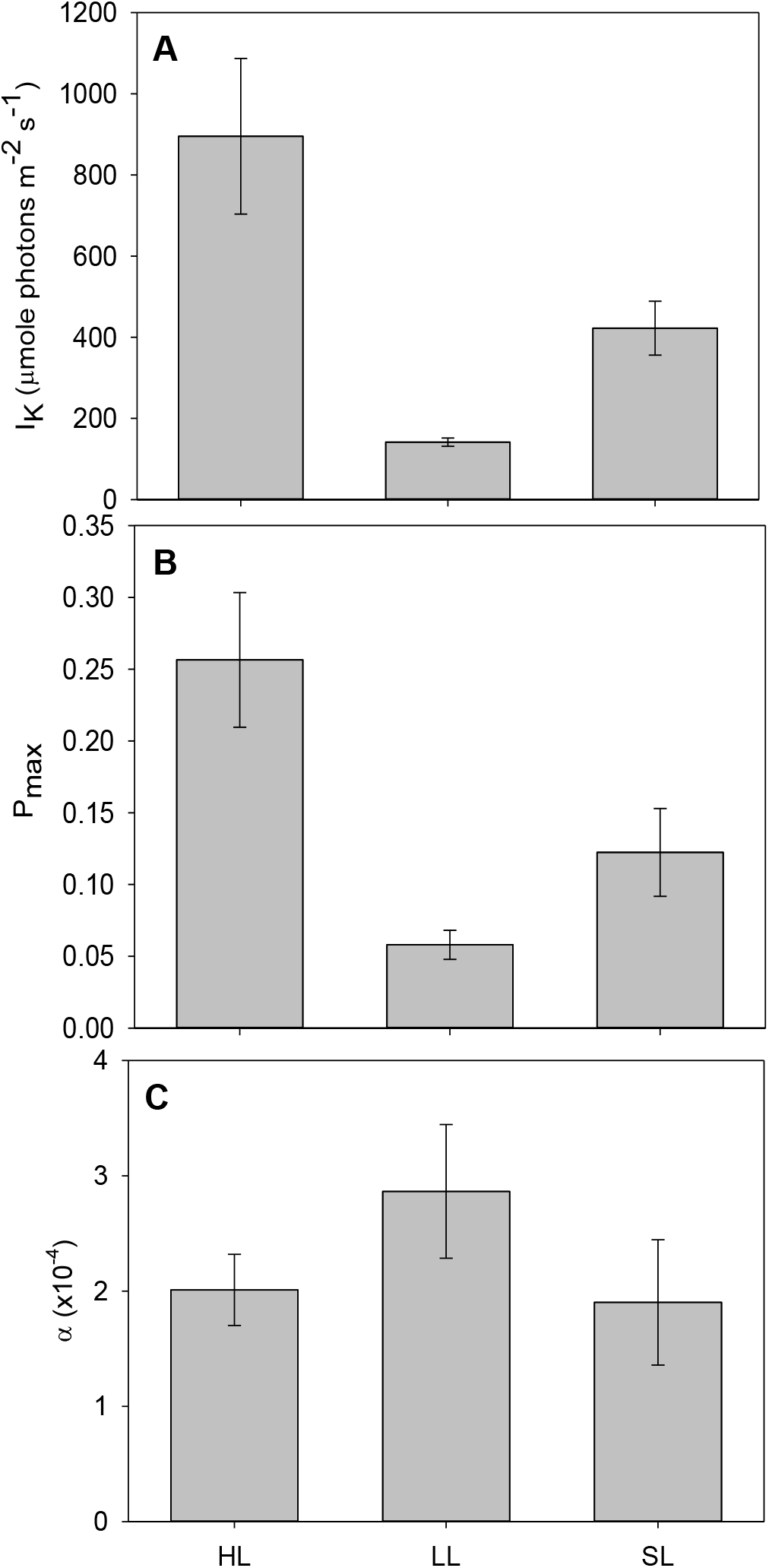
Estimations of photosynthetic performance characteristics (alpha, P_max_, I_K_) from net photosynthesis capacities calculated during an *ex situ* P-I curve. HL and LL correspond to growth with incident light fluxes of 600 μmol photons m^−2^ s^−1^ and 60 μmol photons m^−2^ s^−1^, respectively. SL indicates cultures were grown with sinusoidal daily irradiance in an ePBR photobioreactor. I_K_ (A) represents the calculated minimum light intensity that causes saturation. P_max_ (B) is the maximum predicted net photosynthesis capacity of each acclimated culture. C) Alpha is indicative of the maximum photosynthetic efficiency as a function of oxygen evolution per photon. Error bars represent standard deviation (n=4).

Gross oxygen evolution rates were lower in SL-adapted cells than HL-adapted cells at all light intensities. Oxygen consumption rates in SL cultures were similar to HL, showing minimal changes between 0-470 μmol photons m^−2^ s^−1^, but increasing at each AL interval after this. Net photosynthesis rates were not significantly different from LL-grown cultures throughout the P-I experiment.

Estimates of the maximal rate of photosynthesis suggests the pattern of HL<SL<LL (Fig 5), but cells from all conditions were all able to harvest low light fluxes at equal efficiencies (alpha parameter; Tukey’s test, p≥0.052; Fig 5).

### Estimating in situ photosynthetic capacities in ePBRs

We have previously modelled the light conditions experienced by individual cells in an ePBR system (see Andersson *et al*., 2019 for modelling details). We translated the output of this single-cell model at solar noon to a shortened script for employment on a Dual-PAM 100 coupled with MIMS (Fig. S5). Estimations of PSII excitation pressure (1-qL) showed effective redox balancing of primary electron acceptors during rapid fluctuations in light (Fig. 6A). The highest average 1-qL value observed during these fluctuations was 0.56, suggesting that 44% of acceptors are still in an oxidized state during rapid 0-2,000 μmol photons m^−2^ s^−1^ light transitions. Using a brief period of far-red light after illumination the dark measured qL (qLd) average was 0.97, which infers that only 3% of PSII reaction centers are permanently damaged and required a newly synthesized D1 protein. During these realistic solar noon light fluctuations, NPQ was not significantly employed as a mode of excess energy dissipation, and measured values only reached a maximum of ~0.2 (Fig 6B). MIMS measured oxygen consumption rates showed that gross oxygen consumption is ~90% of gross oxygen production, greatly reducing net oxygen production (Fig 6C).

**Figure 6.**
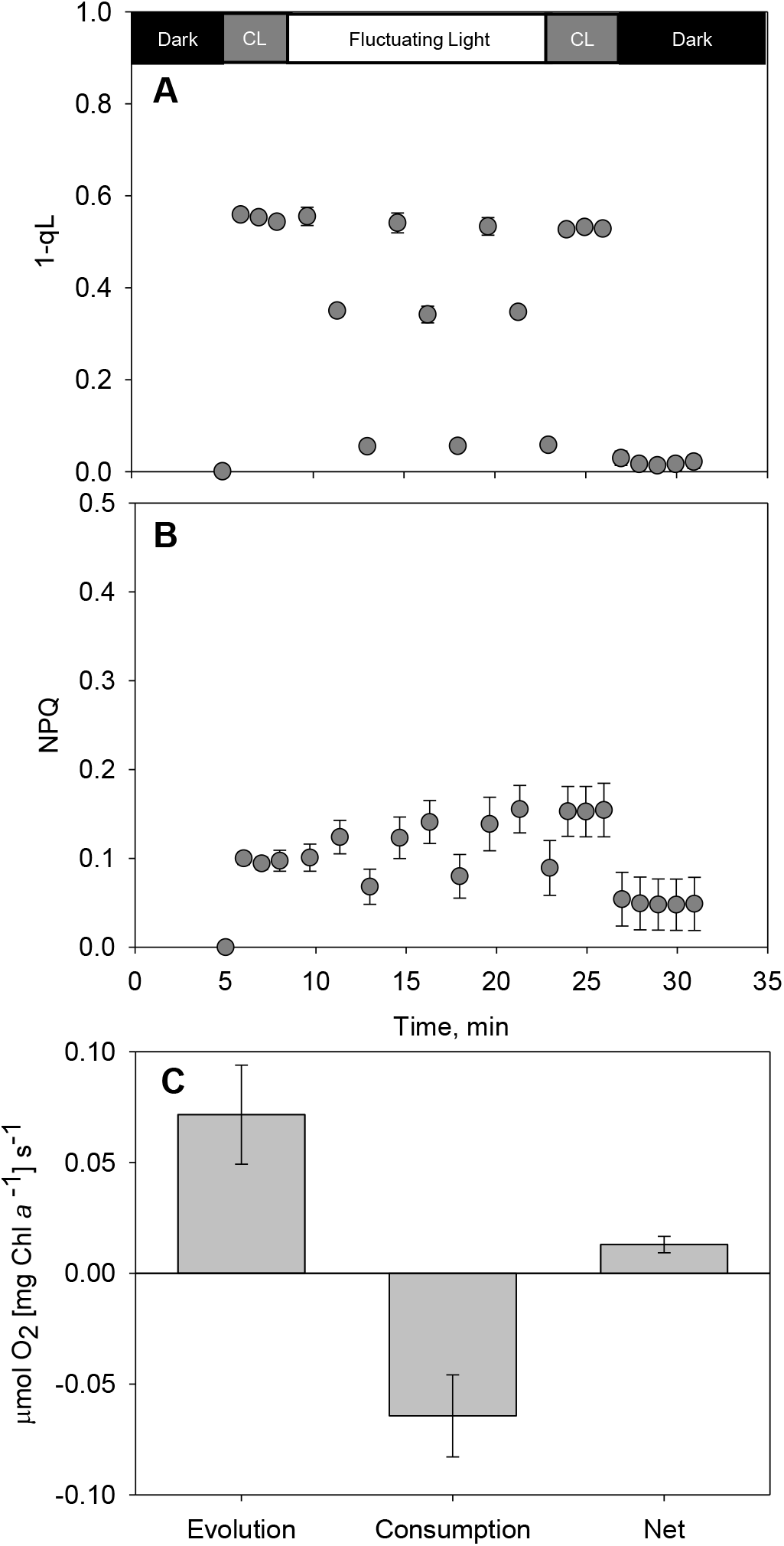
NPQ, Q_A_ redox state (1-qL) and oxygen consumption, evolution and net photosynthesis calculated during *ex situ* experiments replicating the *in situ* cell specific light environment in a benchtop photobioreactor. Chl *a* fluorescence-based estimates of NPQ (A), 1-qL (B) parameters during an *ex situ* fluctuating light experiment that models the cell-specific light environment of ePBR grown cultures at solar noon (see SI for incident light data). Gross oxygen evolution, consumption, and net photosynthesis (C) were measured simultaneously. 2 μg chl *a* was loaded in 1.4 ml total cuvette volume. Error bars represent standard deviation (n=4). CL indicates constant light of 285 μmol photons m^−2^ s^−1^.

### Dynamics of PSII D1 protein repair

500 μg ml^−1^ of the antibiotic lincomycin was applied to HL-acclimated cultures 20 min prior to dawn (ZT -0.33 h) to inhibit chloroplast encoded D1 protein repair in order to distinguish synthesis rates from degradation rates (Key *et al*., 2010; Townsend *et al*., 2018). Fig 7A shows that D1 protein concentrations have visibly declined by ZT 30 min, with an absence of detectible protein by 2hr. Fv/Fm is undermined in lincomycin-treated cultures 15-30 min after dawn (Fig 7B) compared to untreated cultures. Analysis of the decline in Fv/Fm in lincomycin-treated cultures shows that by 42 and 210 minutes of 600 μmol photons m^−2^ s^−1^ light, 50% and 99%< of PSII are damaged respectively (Fig 7B). Analysis of D1 protein Western blots shows that intact D1 protein concentrations starts decreasing at the same rates of Fv/Fm decline (Fig 7C).

**Figure 7.**
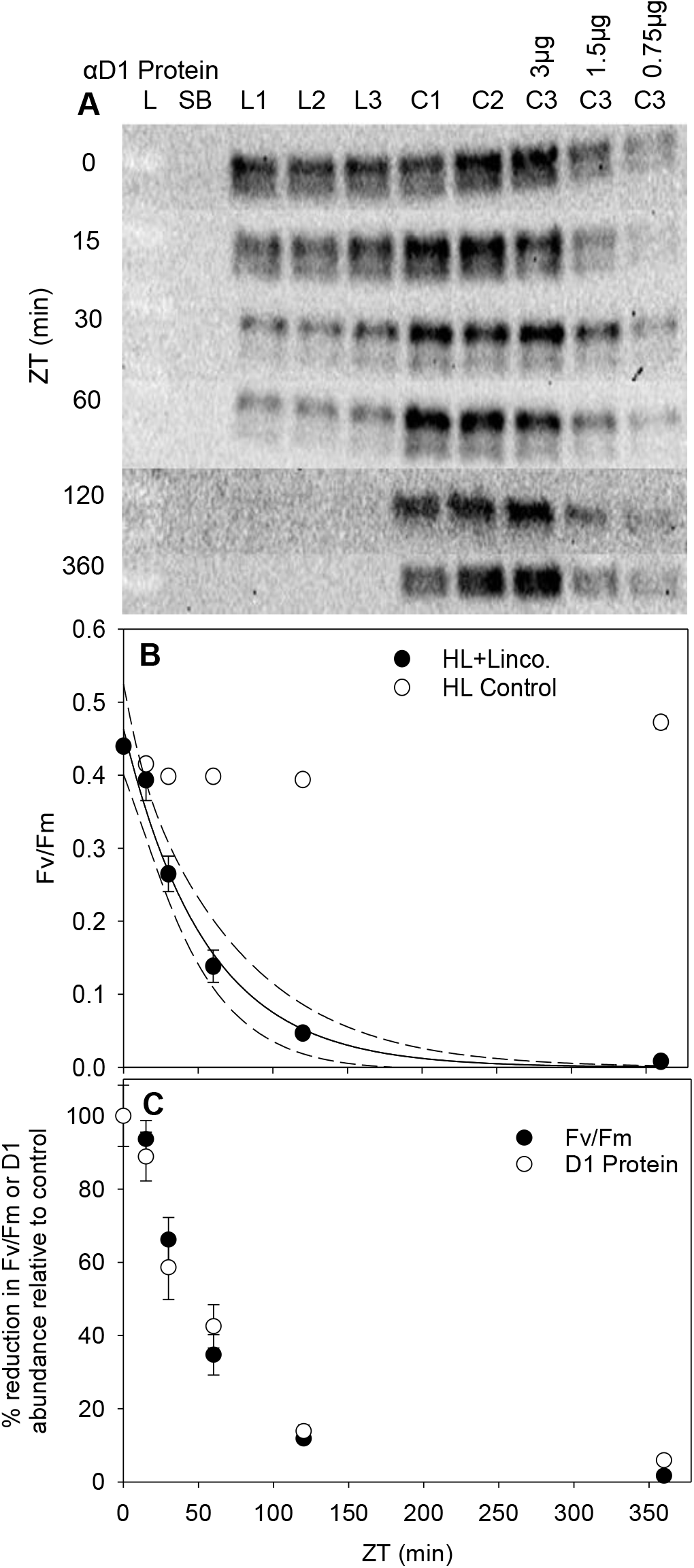
Estimation of D1 protein repair using lincomycin additions to high-light (HL) acclimated cultures at dawn. SDS-Page Western blot analysis of D1 protein using 3 ug of total protein and immunoblot detection of PsbA1 at various time intervals post-dawn (A). C1-C3 are independent cultures exposed to HL (600 μmol photons m^−2^ s^−1^). L1-L3 were treated as controls but with 500 μg ml^−1^ lincomycin added 15 min prior to dawn (ZT -15 min). Error bars represent standard deviation (n=3). L indicates the protein ladder and SB, a sample buffer blank lane. (B) Fv/Fm measured after 30 min dark adaptation. (C) Relative decreases in the abundance of D1 protein abundance and Fv/Fm in lincomycin treated cultures vs. controls. Error bars represent standard deviation (n=3).

## DISCUSSION

### D. armatus dynamically regulates pigment concentrations and avoids photoinhibition

To minimize the overexcitation of photosystems algae can accumulate less pigments per cell, decrease cell size or they can change pigment ratios – which is indicative of a reduction in antenna size. Chlorophyte algae more commonly change ratios and employ state transitions, diatoms reduce pigment concentrations and can functionally dissociate antennae proteins from core photosystem supercomplexes (Cantrell and Peers, 2017; Giovagnetti and Ruban, 2017). Here we show *D. armatus* changes chl *a*/*b* ratio throughout a 24 hr light period, like *Chlamydomonas*, but has significantly higher *a*/*b* ratios. Exponentially growing *D. armatus* has a very small antennae (chl *a*/*b* 6-10) until dusk (ZT 12 h), when chl *a*/*b* ratios decline to <5 (Fig S2). This happens regardless of light conditions. For comparison, *Chlamydomonas* grown under similar HL and LL static light and sinusoidal light conditions had chl *a/b* ratios of 2.76-3.49 (Cantrell and Peers, 2017), and engineered truncated light harvesting antennae protein mutants have chl *a/b* ratios of ~8.1 (Polle *et al*., 2003). It is worthwhile noting the unusual adaptation response in pigment ratio and quantity of *D. armatus* during its exponential growth phase. Chl *b* is almost exclusively located in the light harvesting antennae, thus high chl *a*/*b* ratios indicate small antennae. This suggests that *D. armatus* produces significant amounts of PSII reaction centers (RCII). This is a relatively costly investment, as RCII has more pigments and proteins than antennae complexes and requires continual replacement (discussed further below).

Maximum quantum photochemical yields (Fv/Fm) showed little variation throughout the course of the day in SL and LL grown cultures. HL acclimated cells were the only ones that exhibited photoinhibition at any time during the day, with a decline in PSII yields detected within the first 30 min of exposure to 600 μmol photons m^−2^ s^−1^. This initial decline however was restored to maximum capacity by the next timepoint, indicating an efficient combination of photoprotection and D1 protein repair. We then proceeded to investigate the contribution of various potential mechanisms to maintaining a high ΦPSII in *D. armatus*.

### Diurnal changes in PSII functional antenna size

FIRe measurements offer a lot of photophysiological inferences from PSII chl *a* fluorescence data that allow for estimations of PSII efficiency, size of PSII functional cross section, Fm’ quenching and to distinguish between the size and repair rates of PSII across different algal species (Key *et al*., 2010, Oxborough *et al*., 2012; Jallet *et al*., 2016). We conducted measurements to coincide with solar noon (ZT 6 h) and to test whether midday suppression of PSII occurs, observed as lower Fv/Fm in diatoms, cyanobacteria, and higher plants (Andersson *et al*., 2019; Demmig-Adams *et al*., 1989; Jallet *et al*., 2016). Furthermore, we employed this tool to confirm the inference of a small antennae protein complex, based on the observed high chl *a:b* ratios (Figs 1B and C; Fig S2). These measurements showed that the functional cross section of *D. armatus* does not change drastically during the day and is much smaller (218-238 Å^2^, Fig 2A) than *Phaeodactylum tricornutum* which is ~400 Å^2^ in sinusoidal light conditions (Jallet *et al*., 2016). The induction of NPQ can dramatically increase or decrease the absorption cross section depending on the species-(See Jallet *et al*., 2016; Giovagnetti and Ruban, 2017 for examples). The lack of an observable change in σPSII in the light or dark-adapted states suggested that NPQ is likely not induced in our culture conditions. Indeed, we did not observe *in situ* NPQ (Fig 2B). This agrees with the consensus that NPQ is predominantly located in the major antennae of Chlorophytes and higher plants (Goral *et al*., 2012; Nicol *et al*., 2019), and that *D. armatus* has a constitutively small antennae protein network.

These measurements suggest minimal *in situ* NPQ during the day, but they do not provide conclusive evidence for a lack of photoinhibition. Photochemical quantum yield measurements depend on Fo/Fo’ and Fm/Fm’. NPQ affects both Fo’ and Fm’, and photoinhibition Fo’ (Townsend *et al*., 2018). We utilized the Fo/σPSII parameter to estimate changes in active RCII. The ratio of Fo/σPSII has been used to infer the closure of reaction centers. Oxborough *et al*., (2012) demonstrated a linear positive correlation between Fo/σPSII and active RCII (nmol m^−3^) across Chlorophytes and diatoms. Therefore, the lack of significant changes in Fo/σPSII from ZT 3-9 suggest that the concentration of open reaction centers do not change (Fig 2C). In concert, chlorophyll fluorescence measurements suggest limited midday suppression of light harvesting and effective photoprotective processes in *D. armatus*.

### Desmodesmus does not have a large capacity for NPQ

NPQ is considered to be the major component of photoprotection in eukaryotic algae and plants. Many studies testing NPQ capacities utilize a low light to high light shift, or P-I measurements, which do not occur in mass culture conditions (Andersson *et al*., 2019). Jallet *et al*., (2016) illustrated that a high *ex situ* NPQ capacity, does not mean a large *in situ* NPQ, even when grown under simulated natural light regimes. We wanted to test whether *in situ* conditions here were also insufficient to activate NPQ, or whether NPQ is absent as a major photoprotective mechanism in this species. The P-I experiments illustrated that intense light conditions were not enough to reduce all primary electron carriers in HL and SL-acclimated cultures (measured as 1-qL, Fig 3B). This was correlated with a limited NPQ capacity of <0.4 (Fig 3A). For comparison, SL-grown *P. tricornutum* form the same amount of NPQ at 620 μmol photons m^−2^ s^−1^, as *D. armatus* cells do at 4,000 μmol photons m^−2^ s^−1^ (Jallet *et al*., 2016). *Chlamydomonas reinhardtii* grown and assayed under similar LL, HL and SL conditions had NPQ values of ~0.4, ~0.7 and ~1.0 respectively at 1,850 μmol photons m^−2^ s^−1^ (Cantrell and Peers, 2017). Whereas at 1,750 μmol photons m^−2^ s^−1^ all conditions here yielded NPQ capacities of <0.2. These phenotypes are normally exhibited in mutants of other algal species that do not accumulate antennae proteins and it agrees with NPQ capacities observed in closely related *Scenedesmus* species (Perales-Vela *et al*., 2007; Chalifour and Juneau, 2011; Ioannidis *et al*., 2011).

### Alternative Electron Transport (AET) in Desmodesmus

Previous studies have found that AET can represent a large portion of total electron transport in a wide variety of algae and cyanobacteria (see Radmer and Kok 1976, Kana 1992 for two examples). We estimated the capacity for AET by measuring light dependent oxygen consumption of ^18^O_2_ using MIMS. A large amount of light dependent oxygen consumption significantly reduced the net oxygen production capacity of cells adapted to all light conditions (Fig 4). Furthermore, as is common with photosynthetic organisms, parameters derived to measure photosynthetic efficiency and maximum productivity (Fig 5) reflect acclimation history. The α parameter shows differences between all groups (RM-ANOVA, p=0.042), but differences between HL and LL adapted cells were not significant (Tukey’s test, p=0.084) as in other organisms (Ware *et al*., 2015; Cantrell and Peers, 2017). α is predominantly influenced by the functional antennae of supercomplexes, but LL conditions do not induce large antennae in *D. armatus* during exponential growth phase. Despite the lack of significant acclimation-related changes in α, Pmax and I_K_ parameters change significantly (Fig 5). Thus, it likely reflects differences in electron transport and carbon fixation processes that arise independently of antennae protein accumulation. Indeed, faster growth rates (Table S1), and estimated electron transport rates (Fig S6) add support to these inferences from Pmax and I_K_ parameters.

### AET reduces net photosynthesis in simulated industrial light conditions

Recent modelling of cell specific light environments simulating industrially relevant sinusoidal light conditions have demonstrated that light conditions in photobioreactors are highly dynamic and irregular. Andersson *et al*., (2019) illustrated that cells spend only ~25% of the day in the linear response range (<Phalf-max), and the remainder in either light limiting or excess light conditions, which significantly reduces photosynthetic efficiency compared to growth in the integrated average light intensity. Growth of photosynthetic algae in industrially relevant conditions have suggested that AET plays a more prominent role than NPQ in balancing the overreduction of PSII (Jallet *et al*., 2016; Andersson *et al*., 2019). Testing of NPQ capacity *ex situ* was required to rule it out as a major photoprotective process in this organism, but it might have suggested an exaggerated role for AET *in situ*. Thus, we utilized the light environment previously used by Andersson *et al*. (2019) to assay the utilization of AET vs NPQ in industrially relevant, *in situ* light conditions (ZT 6 h; Fig S5). These assays showed that extreme light fluctuations do not cause an overreduction of Q_A_ and do not induce NPQ (Fig 6A and B). Instead, up to 90% of electrons generated by the splitting of water can be consumed by oxygen consuming reactions downstream of PSII (Fig 6C), and they do not contribute to carbon fixation. Additionally, there is only minor photoinhibition in approximately 3% of RCII (qLd = 0.97). This result suggests that AET engineering could improve photosynthetic efficiency at an industrial scale. Indeed, expression of FLV proteins in tobacco and *Arabidopsis*, and overexpression of FLV proteins in *Synechocystis* sp. PCC6803., can increase carbon assimilation and improve photosynthetic efficiency, particularly in fluctuating light conditions (Hasunuma *et al*., 2014; Yamamoto *et al*., 2016; Gómez *et al*., 2018).

### D1 protein turnover is essential for maintaining a high ΦPSII

HL acclimated cells were chosen to investigate the dynamics of PSII repair, as previous measurements of Fv/Fm variance throughout the course of the day, show that these conditions are enough to cause significant closure of PSII in the first hour after dawn (Fig 1a; ZT 0-1). Lincomycin infiltration is a useful tool as it allows for the separation of photoprotective processes in maintaining a functional PSII, from recovery processes that restore the damaged chloroplast-encoded D1 protein. We observed a decline in Fv/Fm and a ~90% reduction in D1 protein within 2 hours of light exposure in lincomycin treated cells (Figure 8). We estimated that the half-life of the D1 protein in *D. armatus* is 42 min. This is significantly more rapid than has been observed for diatoms (Key *et al*., 2010). D1 protein turnover in spinach illustrated a 30-120 min half-life and the rate was dependent on light acclimation history (Aro *et al*., 1993; Tyystjärvi *et al*., 1994). Because we observed nearly maximal Fv/Fm in HL and in SL conditions, this suggests that *D. armatus* D1 protein capacity repair is fast and extremely efficient. It also suggests that the photoprotective processes described above are not completely adequate for preventing PSII from damage in excess light conditions.

Although beyond the scope of this investigation, the rapid replacement of D1 proteins could be due to a combination of mechanisms. It has been suggested that certain algal species maintain a reserve of surplus PSII subunits to improve the efficiency of degradation and removal of damaged D1 proteins and reinsertion of functional polypeptides (Ni *et al*., 2017). It could also be due to an increase in transcriptional activity of the protease responsible for D1 protein degradation in high light (Hihara *et al*., 2001), or an accumulation of FtsH proteases (Ni *et al*., 2017). An efficient D1 protein turnover, and thus restoration of maximum ΦPSII is a useful trait for biofuel platform species.

## CONCLUSIONS

Higher plants and many algae rely on NPQ to dissipate excess energy accumulated during high or fluctuating light conditions. This work has established that *D. armatus* has three alternate, but effective mechanisms to minimize the accumulation of damaged PSII: efficient D1 protein turnover, a constitutively small antenna and AET. The first two characteristics are desirable for commercial production of algae, with reduction of antennae proteins reflecting a prominent target for bioengineering in order to increase photosynthetic efficiency (Polle *et al*., 2003; Melis, 2009). It was therefore encouraging to observe these properties in an organism already shown to have high productivities in mass culture. This work also adds support to our recent studies on *Synechocystis* sp PCC 6803 (Andersson *et al*., 2019) and *Phaeodactylum tricornutum* (Jallet *et al*., 2016) which suggest that NPQ is minimally employed by cells in light conditions that simulate mass culture, and that AET should be a focus of future algal bioengineering targets to increase photosynthetic productivity. Furthermore, SL-acclimated cultures had the highest maximum theoretical photochemical quantum yields, despite rapid changes in effective light intensities. Square wave light regimes lead to PSII supercomplex arrangements that are not reflective of organisms in nature or growing in sinusoidal light conditions (Ware *et al*., 2015) and this work bolsters the importance of studying photosynthetic organisms in simulated natural light regimes.

## MATERIALS AND METHODS

### Growth conditions and treatments

*Desmodesmus armatus* (SE00107) was grown in modified artificial saltwater media (MASM, pH 8.0]; for complete nutrient composition see Supplemental Table S2). Constant light cultures were grown under 12 h day/night cycles with a custom designed Dynamic Environmental Photosynthetic Imager (DEPI) white LED light bank. High light (HL) conditions correspond to 600 μmol photons m^−2^ s^−1^ and low light (LL) to 60 μmol photons m^−2^ s^−1^ surface intensity measured with a light sensor (Walz ULM-500). Cultures were bubbled with atmospheric air (1 L air L^−1^ culture min^−1^) and grown at 30°C. Samples were collected between ZT 5-7 h after dawn to simulate noon measurements, unless otherwise stated. For cultures grown under a sinusoidal light regime in environmental photobioreactors (ePBRs), the LED lights were programmed to achieve a maximum 2,000 μmol photons m^−2^ s^−1^ surface light intensity at solar noon (see Fig. S1) by the following equation:
Incident Light Flux(t) = A_max_ * sin (2 * π * f * t), where t = s after dawn, A_max_ = 2,000 μmol photons m^−2^ s^−1^, f = 1/(2*Daylength[s]) = 1.16 × 10^−5^ s^−1^.

SL-grown cultures were also bubbled with atmospheric air (1 L air L^−1^ culture min^−1^) and grown at 30°C, with additional stirring (500 rpm). Samples were collected between ZT 5-7 h after dawn to simulate noon measurements, unless otherwise stated.

For lincomycin treatment, control and treated cultures were split in exponential phase. Three flasks were supplemented with 500 μg ml^−1^ lincomycin 20 min before dawn, with three flasks used for controls (Key *et al*., 2010). Biophysical measurements and Western blot analysis were performed on samples collected at 0, 15, 30, 60, 120 and 360 min after dawn (Key *et al*., 2010).

### Cell counts

Cell counts were performed on biological triplicates using a Micromaster microscope (Thermo Scientific) and an improved Bright-Line Neubauer haemocytometer (Sigma-Aldrich). For measurements performed during exponential growth phase, growth rates were calculated as ln[N_t1_/N_t0_]/[t_1_–t_0_] where ln is the natural log of cell concentrations (N_t_) and time (t), between sampling times 0 and 1.

### Absorption spectroscopy for pigment quantification

10 ml of *D. armatus* cultures were harvested and centrifuged at 3,220 x g for 10 min. After discarding the supernatant, the pellet was frozen at −20°C (Schumann *et al*., 2005 and references therein). Pellets were thawed and resuspended in aqueous methanol, vortexed for 10 s, placed in a 72°C bead bath for 30 min being agitated every 10 min, placed in a fridge at 4°C for 15 min and centrifuged at 20,000 x g for 10 min (Sartory and Grobbelaar, 1984). Supernatants were transferred to 1.5 ml semi-micro acrylic cuvettes with 1 cm path lengths. Absorption spectra were scanned from 300-800 nm, with 1-nm increments at a scan rate of 600 nm min^−1^ using a Cary 60 UV–Vis Agilent spectrophotometer. Spectra were zeroed at 800 nm. Chlorophyll concentrations were calculated according to the extinction coefficients presented in Porra *et al*., (1989). Pigment extraction efficiency and comparative chl *b* degradation was compared to methodologies outlined in Porra *et al*., (1990, 2002; Fig S7).

### Chlorophyll PAM fluorescence and membrane-inlet mass spectrometry analysis

A Dual-PAM 100 fluorometer (Walz, Germany), coupled with a quad mass spectrometer (Pfeiffer PrismaPlus QMS200; PTM28612), was employed for simultaneous chlorophyll *a* fluorescence and oxygen evolution and consumption measurements in a custom-made 2 ml glass cuvette with plastic stopper (see Andersson *et al*., 2019 for system details). Cells were harvested during the exponential growth phase, centrifuged (1000 x *g*, 10 min) and resuspended in fresh culture medium to reach a final chl *a* concentration of 1.4 μg chl *a* mL^−1^. Samples were purged with N2 until ^16^O_2_ reached approximately half of atmospheric levels. 2 ml of sample was then transferred to the cuvette and the stopper lowered to the sample meniscus. ^18^O_2_ (Cat # 490474 ALDRICH) was then applied to the sample through the stopper. After ~5 min, once ^18^O_2_ reaches a steady state that is greater than ^16^O_2_, the stopper was lowered to purge any residual gas from the cuvette. Samples were maintained at 30°C during all steps using a Fisher Scientific Isotemp water bath. Samples were illuminated with the measuring light (620 nm) for 30 s to allow for accurate determination of Fo. Saturating pulses of 635 nm light were applied at 10,000 μmol photons m^−2^ s^−1^ for 0.6 s. The rapid light curve function of the DUAL-PAM was used with 2 min actinic light intensity steps of 0, 24, 44, 113, 206, 363, 470, 736, 1141, 1751, 2174, 2709, 3339 and 4087 μmol photons m^−2^ s^−1^. Fv/Fm, NPQ, 1-qL and qLd parameters were calculated as previously described (Kramer *et al*. 2004; Ware *et al*., 2015). Rates of oxygen evolution and consumption were calculated with the following equations (Beckmann K *et al*., 2009):

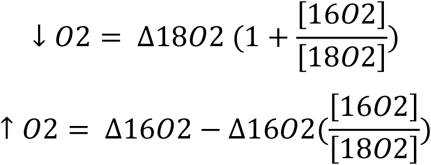

Where [16O_2_] and [18O_2_] reflect the ratios of ^16^O_2_ and ^18^O_2_ at the start of each two-minute actinic light illumination step. Oxygen concentrations were normalized to argon concentrations, chosen for similar solubility as O_2_, to account for abiotic variance due to the MIMS gas consumption (Andersson *et al*., 2019). Ion current slopes were calibrated based on oxygen values for saturated and anoxic media values. Oxygen saturation was achieved by shaking 5-10 ml of media in a flask for 1-2 min. Anoxia was achieved by the addition of sodium dithionite to media. Values were normalized to chl *a* concentration. The minimum saturating light intensity (IK), maximum photosynthetic rate (Pmax) and the light limited slope (α) was calculated by fitting net photosynthetic rates (μmol O_2_ L^−1^ mg chl *a*^−1^ s^−1^) vs irradiance (μmol photons m^−2^ s^−1^) to the empirical functions of Platt *et al*., (1980) using the fitting analysis protocol of Ritchie, (2008).

### Estimations of NPQ and AET in simulated light environments

We utilized the cell-specific light environment calculated for ePBRs as described by Andersson *et al*., (2019) to estimate what photoprotective processes are active in rapidly fluctuating light. 5 min of darkness was used to measure respiration, prior to constant light of 285 μmol photons m^−2^ s^−1^, which was the integrated photon flux intensity at solar noon. This was followed by three 5 min repeats of a rapidly flucuating light script with saturating pulses applied every 100 sec in order to measure Fo’ and Fm’ during the illumination procedure. 5 min of constant light of 285 μmol photons m^−2^ s^−1^ was applied after the dynamic AL illumination, prior to 5 min of darkness. Please see Andersson *et al*. (2019) for details associated with light quality and quantity. PAM fluorescence parameters and MIMS analysis were performed as described above.

### Chlorophyll fluorescence induction and relaxation (FIRe)

Cells were collected at mid-morning, noon and mid-afternoon, corresponding to 3-3.5, 6-6.5 and 9-9.5 h after dawn (Jallet *et al*., 2016). Two times 200 μL of sample were separately added to 1.8 ml of 30°C preheated MASM. One sample was immediately transferred to a sample cuvette for fluorescence analysis, with the other sample being dark adapted for 15 min. A Satlantic FIRe fluorometer was used to measure ΦPSII or Fv/Fm (in light and dark-adapted samples respectively), with 450 nm excitation and 678 nm measured emission (Kolber *et al*., 1998; Jallet *et al*., 2016). σPSII was calculated using Fireworx (https://sourceforge.net/projects/fireworx/) from raw fluorescence data. The Fo/σPSII parameter was subsequently calculated according to Oxborough *et al*., (2012).

### Protein extraction, quantification and visualization

10-30 ml of culture was collected, centrifugated at 3,220 x *g* for 30 min at 4°C, with the resulting pellet resuspended in 1 ml 18,2 MΩ water and frozen at −80°C until subsequent analysis was performed. 100 μl of sample was added to 100 μl of denaturing sample buffer (final concentration 125 mM Tris HCl buffer [pH 6.8], 20% (v/v) glycerol, 4% (w/v) SDS and 0.0025% (w/v) Bromophenol Blue). Homogenization was achieved by vortexing for 30 min with 2 stainless steel beads. Samples were centrifugated for 10 min, 4°C and 20,000 x *g*. The supernatant was heat-treated for 5 min at 65°C (Jallet *et al*., 2016) and then allowed to cool to room temperature. An aliquot was used for the quantification of extracted protein followed using bovine serum standards and Pierce BCA Protein Assay kit, with the remainder being frozen at −80°C until further analysis.

### SDS-page gel electrophoresis and immunoblotting

Once protein stock solutions reached room temperature, an aliquot was added to sample buffer to make 35 μl of 3mg/ml total protein. 25 μl was loaded into a Novex 10–20% Tris-Glycine Gel. A XCell SureLock Mini-Cell System (Thermofisher) was used for gel electrophoresis according to the manufacturer’s directions. 5 μl of protein ladder (ThermoFisher #26617) was used for migration control, with 15 μl of sample buffer used as a control for antibody specificity. Migration was carried out in a Tris-Glycine SDS Running Buffer (25 mM Tris Base [pH 8.3], 192 mM Glycine and 0.1% (w/v) SDS). Proteins were transferred to Invitrolon PVDF Membranes in a transfer buffer (12 mM Tris [pH 8.3], 96 mM glycine and 20% (v/v) methanol) in a XCell II Blot Module. Membranes were blocked overnight at 4°C in Tris-buffered saline (TBS, 50 mM Tris-Cl [pH 7.5], 150 mM NaCl) and 5% w/v skim milk. All the following steps were carried out at room temperature and under constant agitation. Membranes were challenged for 1 h with a PSII D1 protein antibody (Agrisera, #AS0584) diluted 1:10,000 in TBS with 0.5% w/v skim milk correspondingly. Three 5-min washes in TBS with 0.5% w/v skim milk followed. 1:50,000 dilution of secondary antibody (horseradish peroxidase conjugated donkey anti-rabbit, Pierce #31458) in TBS with 0.5% w/v skim milk challenging was carried out for 1 h. Membranes were washed for four 5-min cycles TBS with 0.5% w/v skim milk, and one 5-min wash in TBS, before application of SuperSignal West Femto Maximum Sensitivity Substrate (ThermoScientific) for visualization (Jallet *et al*., 2016). Total protein was visualized using SYPRO-Ruby according to the manufacturer’s instructions (Thermofisher). Band densities were quantified using ImageJ software, version 1.48 (http://imageJ.nih.gov/ij/). Relative abundances were calculated by comparing band densities to the band detected at ZT = 0 h.

### Low temperature fluorescence spectroscopy

HL grown samples were induced into State II by incubating 10 ml of culture with 5 μM FCCP in the dark for 25 min. State I was achieved by placing 10 ml of culture on a shaker in the presence of only far-red light for 1 h. Samples were collected, placed in a Horiba sample holder (Part #5500031800) and immediately frozen in liquid nitrogen. Measurements were performed at 77K using a Horiba Jobin-Yvon Fluorolog-3 spectrofluorometer and Horiba Liquid Nitrogen Dewar (FL-1013). Excitation was provided at 435 nm, with 1 nm fluorescence spectral resolution with a spectral bandwidth of 5 nm. The baseline was subtracted at 750 nm and normalized to 682 nm.

### Statistical analysis

Statistical differences were tested by a repeated measures one-way analysis of variance test (RM-ANOVA). Tukey’s post-hoc test was utilized to test differences between groups. A paired Student’s t-test was used to compare temporally separated measurements on the same samples in Fig 2. Statistical tests were all performed in SigmaPlot 13.0 (Systat Software, Inc., Chicago, USA). All values reported are means from 3-6 biological replicates ± the standard deviation.

## SUPPLEMENTAL DATA

These materials are available:

Supplemental Figure S1. Comparison of culture growth regimes.

Supplemental Figure S2. Chl *a/b* ratios of cultures based on pigment extractions performed in Fig 1.

Supplemental Figure S3. State-transition capacity of *D. armatus* using PAM fluorescence and 77K fluorescence spectroscopy.

Supplemental Figure S4. UV-B induced NPQ of sinusoidal light acclimated cultures. Supplemental Figure S5. Depiction of the fluctuating light regime to estimate *in situ* NPQ and AET of sinusoidal light acclimated cultures.

Supplemental Figure S6. Estimated electron transport rates from an *ex situ* P-I curve. Supplemental Figure S7. Comparison of solvent extraction protocols in *D. armatus* and *C. reinhardtii*.

Supplemental Table S1. Cell counts, Fv/Fm and chl *a/b* ratios of high light, low light and sinusoidal light acclimated cultures.

Supplemental Table S2. Nutrient composition of modified artificial seawater media (MASM). Supplemental Table S3. Maximum photochemical quantum yield (Fv/Fm) parameters for cultures with different acclimation histories and at different timepoints.

Supplemental Table S4. Chlorophyll *a* (pg) per cell parameters extracted in aqueous methanol.

Supplemental Table S5. Chlorophyll *b* (pg) per cell parameters extracted in aqueous methanol.

Supplemental Table S6. FIRe parameters calculated at ZT 3-9 h for cultures acclimated to different light conditions.

Supplemental Table S7. 1-qL and NPQ parameters calculated from *ex situ* P-I curves. Supplemental Table S8. Oxygen evolution, consumption and net photosynthesis parameters calculated from *ex situ* simultaneous MIMS and P-I curve induction experiments.

Supplemental Table S9. Alpha, Pmax and I_K_ parameters calculated from net photosynthesis capacities measured during *ex situ* simultaneous MIMS and P-I curve induction experiments.

## Abbreviations

AET: alternative electron transfer
RM-ANOVA: repeated measures analysis of variance
chl *a*: chlorophyll *a*
chl *b*: chlorophyll *b*
ePBR: phenometrics environmental photobioreactor
FIRe: fluorescence induction and relaxation
Fv/Fm: maximum quantum yield of photosystem II
h: hour
NPQ: non-photochemical quenching of chl *a* fluorescence
PAM: pulse amplitude modulated
PAR: photosynthetically active radiation
P-I: photosynthesis versus irradiance
PSII: photosystem II
ΦPSII: quantum yield of photosystem II
ZT: Zeitgeber time (hours past dawn)
σPSII: PSII functional absorption cross section.

